# Rapid and Inexpensive Preparation of Genome-Wide Nucleosome Footprints from Model and Non-Model Organisms

**DOI:** 10.1101/870659

**Authors:** Laura E McKnight, Johnathan G Crandall, Thomas B Bailey, Orion GB Banks, Kona N Orlandi, Vi N Truong, Grace L Waddell, Elizabeth T Wiles, Drake A Donovan, Scott D Hansen, Eric U Selker, Jeffrey N McKnight

## Abstract

Eukaryotic DNA is packaged into nucleosomes, the smallest repeating unit of chromatin. The positions of nucleosomes determine the relative accessibility of genomic DNA. Several protocols exist for mapping nucleosome positions in eukaryotic genomes in order to study the relationship between chromatin structure and DNA-dependent processes. These nucleosome mapping protocols can be laborious and, at minimum, require two to three days to isolate nucleosome-protected DNA fragments. We have developed a streamlined protocol for mapping nucleosomes from *S. cerevisiae* liquid culture or from patches on solid agar. This method isolates nucleosome-sized footprints in three hours using 1.5 ml tubes with minimal chemical waste. We validate that these footprints match those produced by previously published methods and we demonstrate that our protocol works for *N. crassa* and *S. pombe*. A slightly modified protocol can be used for isolation of nucleosome-protected DNA fragments from a variety of wild fungal specimens thereby providing a simple, easily multiplexed and unified strategy to map nucleosome positions in model and non-model fungi. Finally, we demonstrate recovery of nucleosome footprints from the diploid myeloid leukemia cell line PLB-985 in less than three hours using an abbreviated version of the same protocol. With reduced volume and incubation times and a streamlined workflow, the described method should be compatible with high-throughput, automated creation of MNase-seq libraries. We believe this simple validated method for rapidly producing sequencing-ready nucleosome footprints from a variety of organisms will make nucleosome mapping studies widely accessible to researchers globally.

## Introduction

Eukaryotic genomes are packaged into chromatin, where octamers of histone proteins are wrapped by ~146 base pairs of DNA to form repeating units known as nucleosomes (Kornberg 1974; Kornberg and Thomas 1974; Luger *et al.* 1997). These nucleosomes are distributed nonrandomly throughout the genome and regulate access of underlying DNA to processes that require DNA association, such as transcription, replication, DNA repair and transcription factor binding (MacAlpine and Almouzni 2013; Venkatesh and Workman 2015; Hauer and Gasser 2017). Because the positions of nucleosomes play an integral role in regulating DNA-dependent processes, techniques to determine the positions of nucleosomes on the genome are employed widely across organisms (Lee *et al.* 2007; Mavrich *et al.* 2008; Valouev *et al.* 2008; Valouev *et al.* 2011).

One common strategy to map nucleosome footprints uses micrococcal nuclease (MNase), which preferentially digests DNA between nucleosomes to leave behind nucleosome-protected stretches of DNA. This has more recently been coupled to high-throughput sequencing to obtain genome-wide nucleosome positioning profiles for many eukaryotic organisms (Henikoff *et al.* 2011; Cui and Zhao 2012). Since there is wide interest in understanding how proper nucleosome positions are established and altered within an organism in various environmental conditions, nucleosome mapping protocols continue to be used to dissect fundamental mechanisms in chromatin biology.

Sequencing of nucleosome-protected DNA fragments has uncovered a detailed understanding of mechanisms leading to regular nucleosome positioning in yeast and other organisms (Lantermann *et al.* 2010; Gkikopoulos *et al.* 2011; Zhang *et al.* 2011; Pointner *et al.* 2012; Krietenstein *et al.* 2016; Wiechens *et al.* 2016; Baldi *et al.* 2018). More recently, sophisticated technology has been adopted to map nucleosomes with higher precision (Brogaard *et al.* 2012; Chereji *et al.* 2018) or to quantitatively gauge true nucleosome occupancy (Chereji *et al.* 2019; Oberbeckmann *et al.* 2019). Nevertheless, there is still a significant need for simple and robust measurement of nucleosome positions across genomes. For this reason, various protocols for mapping nucleosome positions have been established in a wide range of organisms. These protocols are notably different not only between organisms, but can deviate significantly even within a single organism with highly variable cell number input, sample and reagent volumes, DNA purification methods and time commitments (Table 1). These differences make it challenging to identify an appropriate nucleosome mapping protocol for any specific organism and make it difficult to create a new mapping strategy for organisms that lack established protocols.

**Table 1.** Comparison of different protocols for isolating nucleosome footprints.

|  | Starting Material | Formaldehyde Volume | DNA Purification Method | Time Required to Isolate Nucleosome Footprints<sup>a</sup> |
| --- | --- | --- | --- | --- |
| ***S. cerevisiae* (log growth)** |  |  |  |  |
| Rodriguez et al 2014 | 200 ml liquid culture | 5.5 ml | Phenol/chloroform extraction and ethanol precipitation | 2 days |
| Kubik et al 2015 | 50 ml liquid culture | 1.4 ml | Phenol/chloroform extraction and ethanol precipitation | ~1-1.5 days |
| Cole et al 2012 | 250 ml liquid culture | None | Phenol/chloroform extraction and ethanol precipitation | ~1.5 days |
| Rapid Protocol | 25 ml liquid culture | 650 ul | MinElute column | 3 hours |
| ***S. cerevisiae* (quiescent)** |  |  |  |  |
| McKnight et al 2015 | Purified quiescent cells from 25 ml culture | 650 ul | Phenol/chloroform extraction and ethanol precipitation | 2 days |
| Rapid Protocol | Purified quiescent cells from 25 ml culture | 650 ul | MinElute column | ~4 hours |
| ***S. cerevisiae* (agar plate)** |  |  |  |  |
| Rapid Protocol<sup>b</sup> | Patch on agar plate (~10<sup>8</sup> cells) | 27 ul | MinElute column | 3 hours |
| ***S. pombe*** |  |  |  |  |
| Cam and Whitehall | 100 ml liquid culture | 2.7 ml | Phenol/chloroform extraction and ethanol precipitation | 2 days |
| Rapid Protocol | 25 ml liquid culture | 650 ul | MinElute column | 3 hours |

**Table 1 (cont)**
| ***N. crassa*** |  |  |  |  |
| --- | --- | --- | --- | --- |
| Klocko et al 2019 | Nuclei isolated from 500 ml conidia | None | MinElute column | 2 days |
| Seymour et al 2016 | Chromatin fraction from 50 ml conidia | 275 ul | Phenol/chloroform extraction and ethanol precipitation | 3 days |
| Rapid Protocol | 50 ml conidia | 1.4ml | MinElute column | 3 hours |
| **Mammalian Cells** |  |  |  |  |
| Schwartz et al 2019 | HeLa cells (150mm plate, 70-80% confluence) | None | Ethanol precipitation | ~2 days |
| Ramani et al 2019 | Mouse embryonic stem cells ($2.5 \times 10^6$ cells) | None | Phenol/chloroform extraction and ethanol precipitation | < 1 day |
| Rapid Protocol | $10^6$ human diploid cells (PLB-985) | 27 ul | MinElute column | ~2.5 hours |
| **Wild Mushrooms (*various species*)** |  |  |  |  |
| Rapid Protocol<sup>b</sup> | 50 mg of fruiting body | 27 ul | MinElute column | 3 hours |

<sup>a</sup>Times were estimated based on interpretation of described protocols
<sup>b</sup>No other protocols were identified to isolate nucleosome footprints for these types of samples

Here we have created a unified and streamlined strategy that isolates nucleosome footprints in roughly 3 hours, providing a rapid and validated means to create high-throughput sequencing libraries in a single day. Importantly, the same protocol can be used across model organisms. We show that our protocol works in *S. cerevisiae* (both liquid culture and solid patches), *S. pombe*, *N. crassa*, and human cells. In addition, we tested the protocol on multiple wild mushrooms and efficiently recovered nucleosome footprints, suggesting this protocol can likely be adapted as a starting point to map nucleosome positions in model and non-model eukaryotes.

## Results

### Streamlined Protocol for Rapid Recovery of Nucleosome Footprints from S. cerevisiae

The isolation of nucleosome-protected DNA is typically achieved through a similar procedure across all established protocols. For yeast, cells are generally crosslinked with formaldehyde to maintain nucleosome positions throughout the subsequent steps, though it has been debated whether these crosslinks can efficiently trap nucleosome positions without introducing artifacts (Henikoff *et al.* 2011; Cole *et al.* 2012). The cell walls are broken and nuclei are permeabilized to allow micrococcal nuclease to access and digest extranucleosomal DNA.

Cellular RNA is removed, formaldehyde crosslinks are reversed and protein is digested prior to DNA purification. Finally, the residual 3` phosphates (from MNase cleavage activity) are removed prior to genomic library construction. Our protocol preserves all of these standard steps, but optimizes the workflow to achieve this process using small volumes while eliminating an entire day of experiment time (Figure 1A, Table 1). The largest time savings compared to other fungi-specific MNase protocols are 1) reduction of time for proteinase K treatment and formaldehyde crosslink reversal from overnight to 45 minutes and 2) elimination of phenol/chloroform extraction and ethanol precipitation in favor of column-based DNA purification.

**Figure 1.**
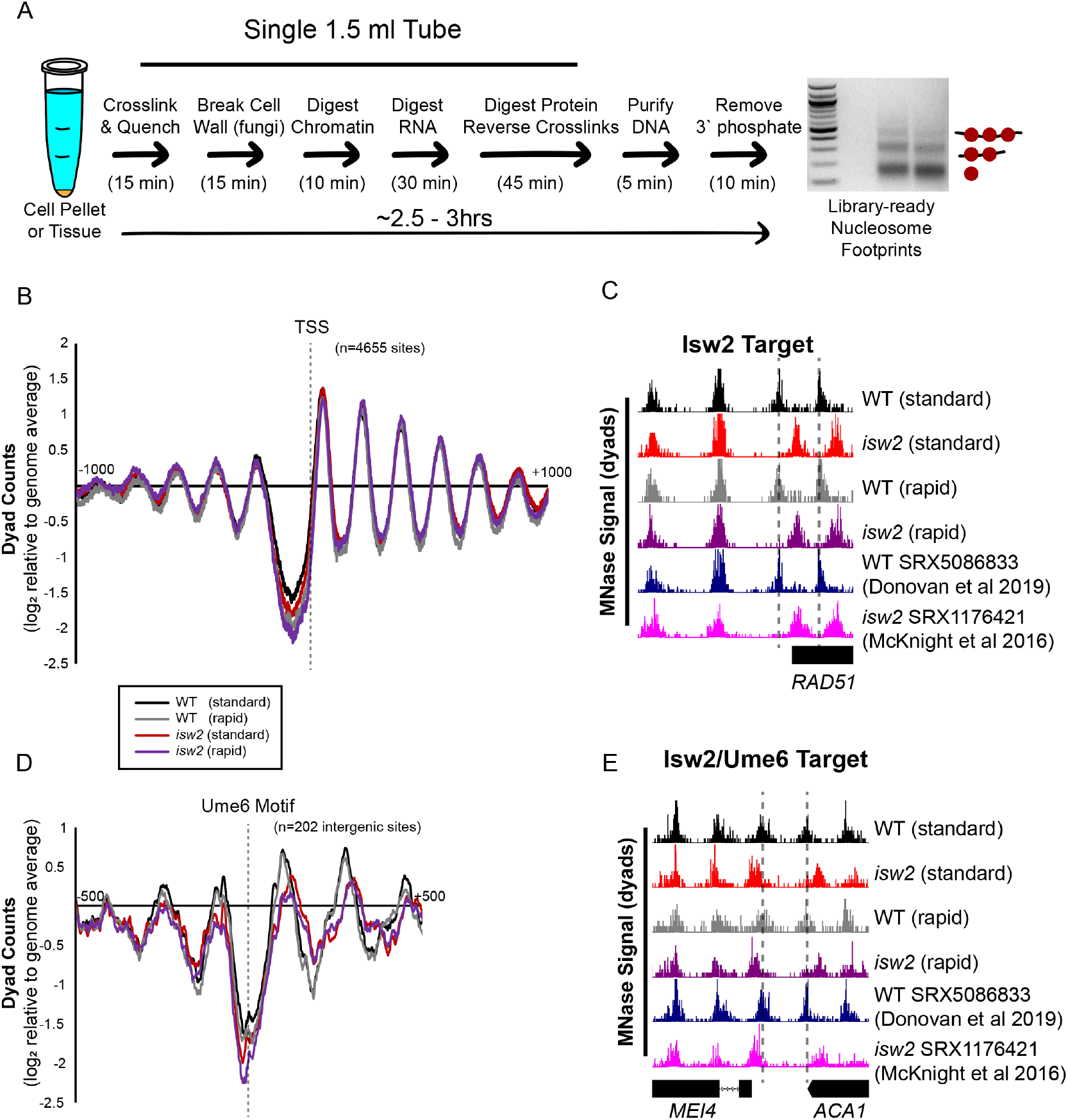
Rapid MNase can Accurately Map Nucleosome Positions in *S. cerevisiae* Cells. (A) Schematic of rapid MNase workflow beginning with a pellet (~200 million cells) of uncrosslinked *S. cerevisiae* cells in a 1.5 ml tube. Cells are crosslinked, spheroplasted, treated with MNase, RNase A, proteinase K while reversing crosslinks, and DNA is purified by spin column prior to phosphatase treatment. A 2% agarose gel showing an example digestion of WT and *isw2* yeast from exponentially-growing cells in YPD is shown (right). (B) Nucleosome dyad signal at 4655 yeast transcription start sites (TSSs) are plotted for WT and *isw2* nucleosomes harvested using a standard (Rodriguez *et al.* 2014) or rapid protocol. (C) Example Genome Browser image showing the standard method and rapid method can map Isw2-directed nucleosome positions similar to published data sets at the *RAD51* locus. Dashed lines indicate Isw2-positioned nucleosomes. (D) Nucleosome dyad signal at 202 intergenic Ume6 target sites showing rapid and standard MNase methods can accurately identify global changes in nucleosome structure at an Isw2 recruitment motif. (E) Genome Browser image showing all methods correctly identify Isw2-positioned nucleosomes at the *MEI4-ACA1* locus, a Ume6 target site. Dashed lines indicate Ume6- and Isw2-positioned nucleosomes.

While the rapid MNase strategy can reproducibly recover well-digested nucleosome ladders, we wished to validate that the recovered nucleosome footprints did not deviate from expected genomic positions. To validate the rapid MNase approach, we created genome-wide MNase-seq libraries and mapped nucleosome positions in a wild type and *isw2* yeast using the new protocol and a previously-published protocol (Rodriguez *et al.* 2014). We chose Isw2-deficient yeast because Isw2 is required for nucleosome shifts at specific target loci, which allows us to determine if our protocol can recapitulate Isw2-specific nucleosome positions at target sites. For all samples, nucleosome organization at transcription start sites (TSSs) displayed the stereotypical structure, with a nucleosome-depleted region flanked by packed nucleosome arrays (Figure 1B). Comparison of nucleosome positions at Isw2 targets showed that nucleosomes were detected at strain-specific but not protocol-specific locations (Figure 1C). Both the rapid protocol and standard protocol recovered strain-specific nucleosome positioning at Ume6 binding sites, a known Isw2-recruitment protein (Goldmark *et al.* 2000), across the genome (Figure 1D,E). In sum, the rapid protocol can accurately map nucleosome positions and capture strain-specific nucleosome footprints.

We next asked whether the rapid MNase protocol could be applied to purified quiescent cells (Allen *et al.* 2006). To test this, we first purified quiescent yeast from 3-day saturated cultures using a Percoll gradient. As previously observed, quiescent yeast required longer treatment with an increased amount of Zymolyase (McKnight *et al.* 2015) to compensate for a highly-fortified cell wall (Li *et al.* 2015). We observed that the rapid protocol with an extended Zymolyase incubation can efficiently and reproducibly recover nucleosome footprints (Figure 2A). Because of the protocol’s simplicity, we suspected that we could similarly recover nucleosome footprints from patches of yeast grown on agar plates, or “colony MNase”. Indeed, nucleosome footprints are readily recovered from solid-grown yeast (Figure 2B), and the captured nucleosome positions accurately reflect nucleosome positioning across the yeast genome (Figure 2C,D). Similar to liquid culture, the rapid colony MNase protocol can also detect Isw2-specific nucleosome positioning events genome-wide (Figure 2E,F).

**Figure 2.**
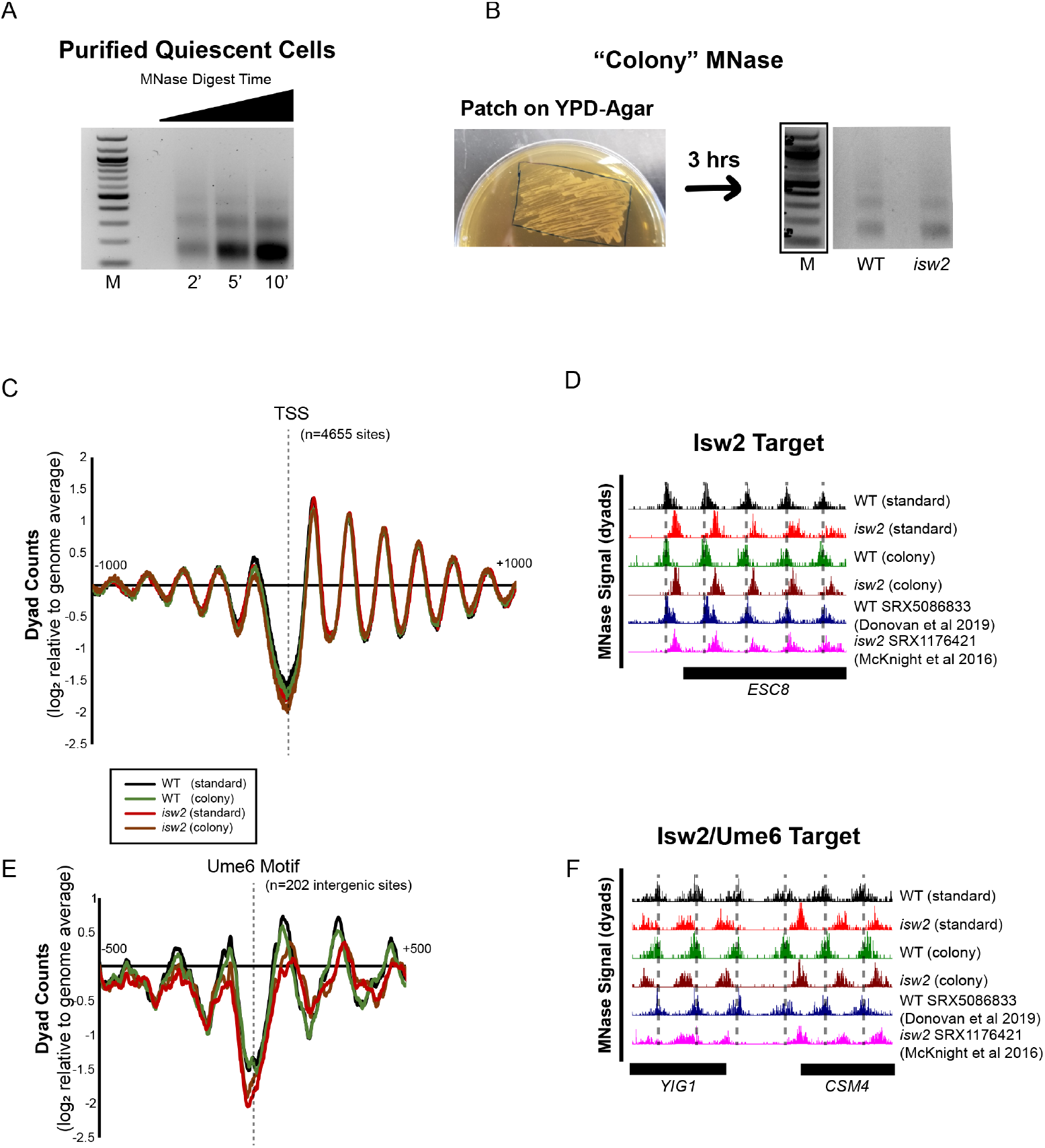
Rapid MNase can Recover Nucleosome Footprints from Isolated Quiescent Cells and Yeast Patches. (A) Representative gel showing nucleosome footprints recovered from purified quiescent cells using the rapid MNase protocol. (B) Representative gel (right) showing nucleosome footprints recovered from a fresh patch of yeast collected from YPD-Agar (left). (C) Nucleosome dyad signal at transcription start sites (TSSs) comparing standard (Rodriguez et al. 2014) and patch-recovered “colony” MNase footprints. (D) Example Genome Browser image showing “colony” MNase footprints can accurately identify Isw2-directed nucleosome positions at the ESC8 locus compared to the standard MNase protocol and published data sets. (E) Nucleosome dyad signal at 202 intergenic Ume6 target sites showing “colony” MNase can accurately identify global changes at an Isw2 recruitment motif. (F) Genome Browser image showing colony MNase can similarly identify Isw2-positioned nucleosomes at the YIG1-CSM4 locus, an example Ume6 target site. Dashed lines indicate Ume6- and Isw2-positioned nucleosomes.

### Rapid Recovery of Nucleosome Footprints from S. pombe and N. crassa

After validating the rapid MNase protocol as a useful and versatile method to map nucleosomes in *S. cerevisiae* we wished to determine if the same protocol could be used for other fungal species. To test this, we first grew *S. pombe* in liquid culture and prepared MNase-seq libraries after using the rapid MNase protocol. Without any protocol modifications, we were able to recover a well-defined and appropriately-digested nucleosome ladder from wild-type *S. pombe* cells (Figure 3A). We compared our sequencing data set with previously-published MNase-seq data sets from *S. pombe* cells (DeGennaro *et al.* 2013; Steglich *et al.* 2015). Genomic nucleosome dyad positions from samples prepared by the rapid MNase protocol were the same as seen previously (Figure 3B). In addition, global nucleosome positioning at *S. pombe* TSSs was nearly identical across data sets indicating that rapid MNase can readily capture global nucleosome positions in *S. pombe* (Figure 3C). We next tried the rapid MNase protocol on crosslinked, germinated *Neurospora crassa* conidia. Similar to both *S. cerevisiae* and *S. pombe* cells, we were readily able to isolate nucleosome footprints from *N. crassa* using the rapid MNase protocol (Figure 3D). Importantly, we verified that the genomic nucleosome positions that we obtained were similar to previously-published *N. crassa* data sets (Seymour *et al.* 2016; Klocko *et al.* 2019) (Figure 3E). Together these data validate the rapid MNase protocol as a reasonable, rapid strategy to map nucleosome positions in *S. pombe* and *N. crassa*, two widely-investigated model fungi.

**Figure 3.**
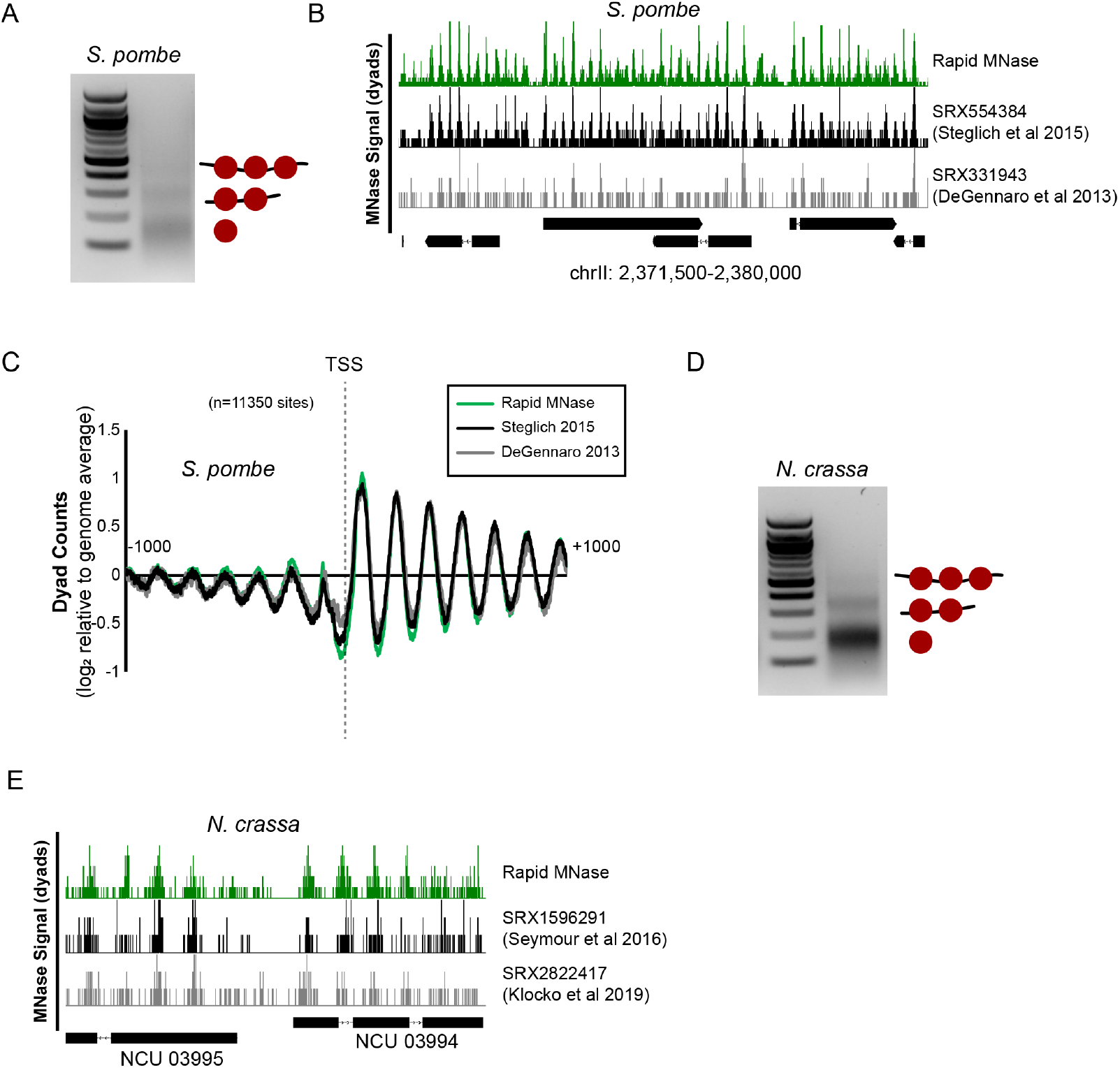
Rapid MNase can Accurately Map Nucleosome Positions in *S. pombe* and *N. crassa*. (A) Agarose gel showing example nucleosome footprints recovered from *S. pombe* using the rapid MNase protocol. (B) Genome Browser image comparing nucleosome dyad positions on *S. pombe* chrII recovered for the rapid MNase protocol (top) and previously-published data sets. (C) Alignment of nucleosome dyads at 11,350 transcription start sites (TSSs) for rapid MNase-recovered nucleosome footprints and previously-published data sets. (D) Agarose gel showing example nucleosome footprints recovered from *N. crassa* using the rapid MNase protocol. (E) Genome Browser image comparing nucleosome dyad positions at the *N. crassa* NCU3995-NCU3994 locus for the rapid MNase protocol (top) and previously-published data sets.

### Recovery of Nucleosome Footprints from Wild Fungal Specimens

After validating our protocol on multiple model fungi, we next asked if we could apply the rapid MNase protocol to non-model fungi. We foraged for local wild mushrooms and subjected them to the rapid MNase workflow. Interestingly, we successfully isolated mononucleosome-sized footprints from all wild mushrooms that we collected (Figure 4A). After optimizing the amount of input mushroom tissue to 50 mg, we could reproducibly obtain similarly-digested nucleosome ladders from multiple mushrooms, including commercially-important chanterelles, in under three hours (Figure 4B, top). Application of the optimized protocol to a newly-acquired specimen led to successful recovery of well-digested chromatin (Figure 4B, bottom) suggesting that nucleosomes can be digested and recovered from many wild mushrooms using this protocol. This successful isolation of nucleosomes from previously-untested organisms suggests that the rapid MNase protocol can be applied across a wide variety of fungal species, and will be a useful and easy-to-implement approach to define MNase protocols in new systems.

**Figure 4.**
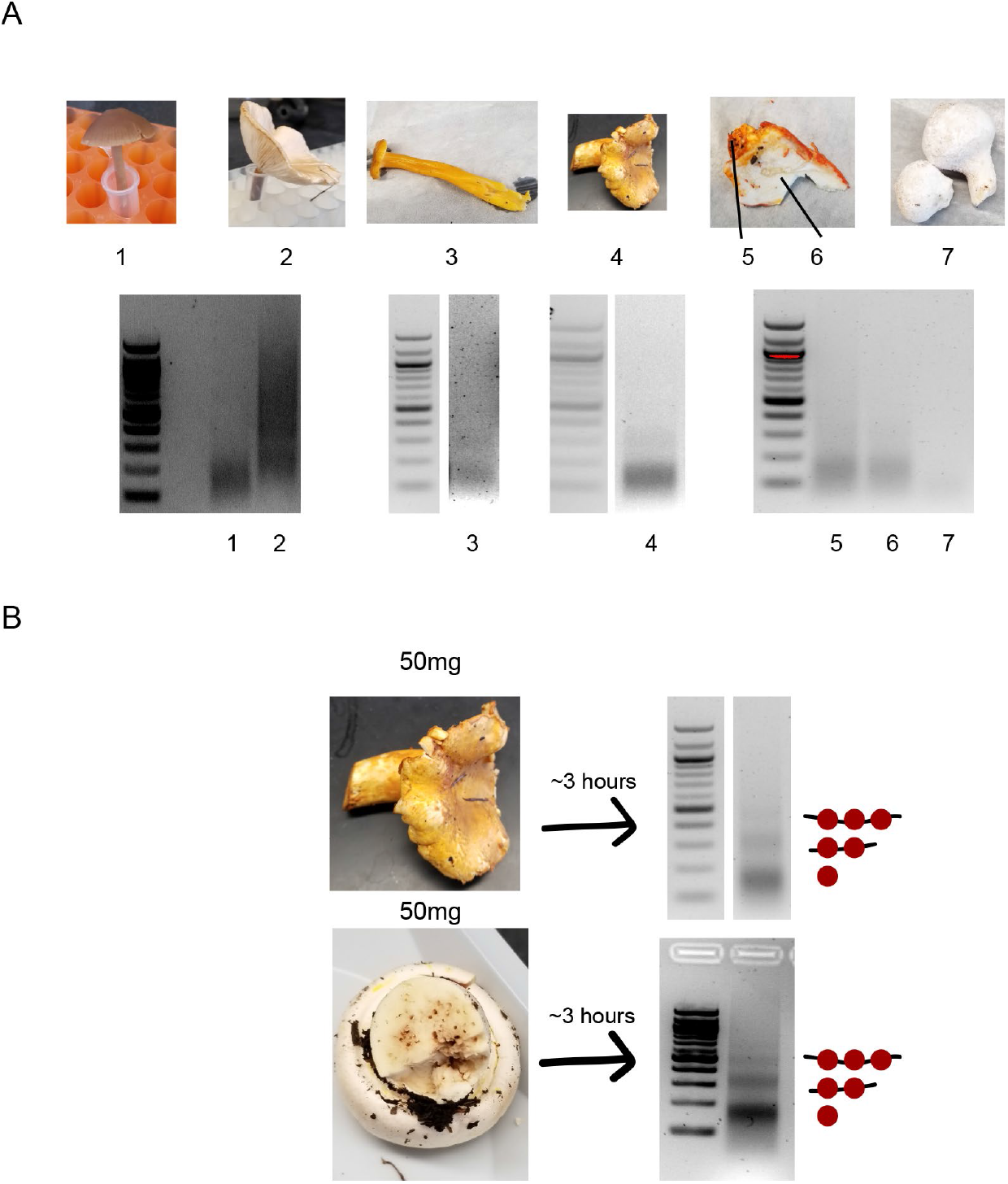
Nucleosome Footprints can be Rapidly Recovered from Wild Mushroom Samples. (A) Images of locally-foraged wild mushrooms that were subjected to the rapid MNase protocol (top). Sample 5-6 consists of a distinct surface fungal specimen (5) growing on a host specimen (6). Recovered nucleosome footprints for corresponding mushrooms are shown (bottom). Speculative identities of these samples are (1) *Panaeolus foenisecii*, (2) unknown, (3) *Craterellus tubaeformis*, (4) *Cantharellus formosus*, (5) *Hypomyces lactifluorum*, (6) *Russula brevines*, (7) *Lycoperdon perlatum*. (B) Optimized rapid MNase for specimen 4 (chanterelle) was achieved using 50 mg of tissue leading to well-digested nucleosome footprints (top). The optimized protocol was performed on 50 mg of a previously-untested specimen leading to well-digested nucleosome footprints (bottom). Speculative identity of these samples are *Cantharellus formosus* (top) and *Agricus xanthodermus* (bottom).

### Recovery of Nucleosome Footprints from the PLB-985 Leukemia Cell Line

Although the rapid MNase protocol was initially designed to expedite the recovery of nucleosomes from *S. cerevisiae*, our success in obtaining nucleosome footprints from multiple fungal species suggested that the protocol may be broadly applicable in other organisms. We tested whether we could use the rapid MNase protocol (without cell wall digestion) to isolate nucleosome footprints from PLB-985 cells, a diploid myeloid leukemia cell line. With a formaldehyde crosslinking step, reproducible nucleosome footprints were robustly recovered in less than 2.5 hours (Figure 5A). If crosslinking is omitted, since it is potentially less likely that nucleosome positions will change in the extremely short duration of the protocol, it is possible to recover library-ready nucleosome footprints in less than 1 hour (Figure 5B). Although previous MNase protocols in mammals could be performed in less than 1 day (Ramani et al 2019), the fact that our rapid protocol described herein can be used across organisms makes it a promising and standardized alternative to previously-published protocols across organisms.

**Figure 5.**
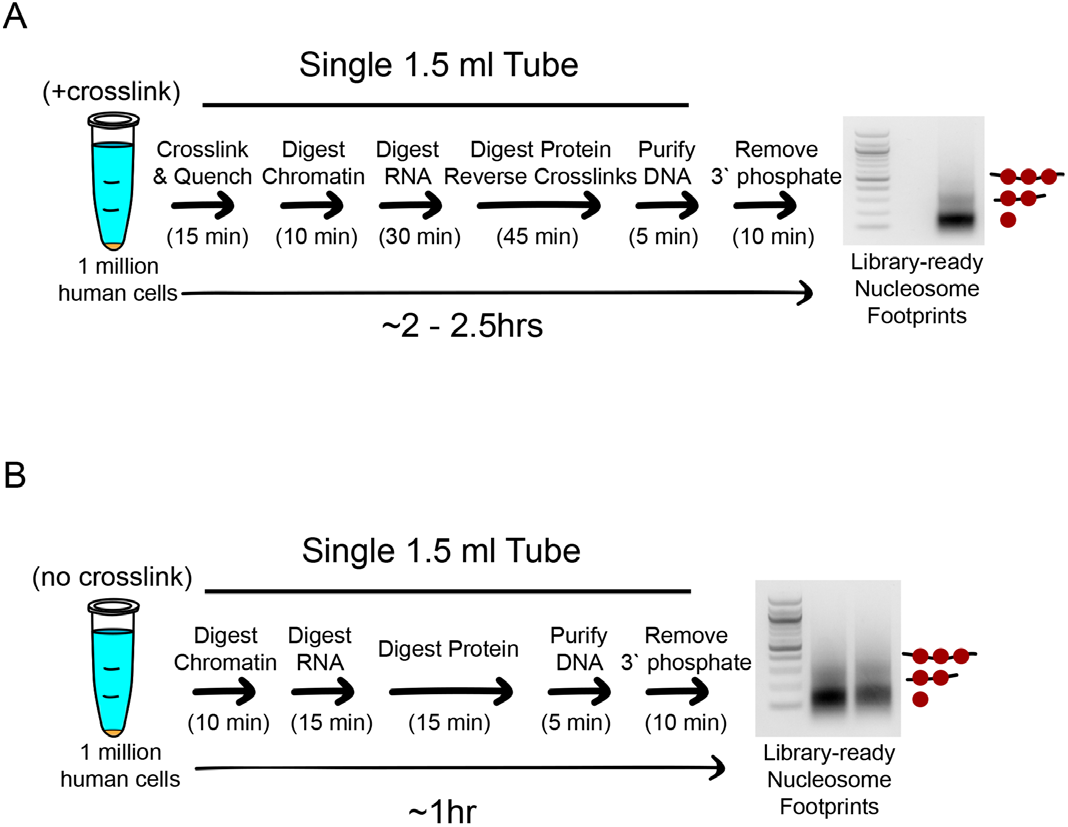
The Rapid MNase Protocol can be Performed on Human Cells with or without Crosslinking. (A) Cartoon schematic showing rapid MNase protocol and associated nucleosome footprints for 1 million human cells from the diploid myeloid leukemia cell line PLB-985. The protocol is identical to that in Figure 1A except there is no Zymolyase treatment. (B) Cartoon schematic showing rapid MNase protocol for PLB-985 cells and associated nucleosome footprints when the crosslinking and crosslink reversal steps are omitted and other steps are shortened. The crosslink-free protocol can provide nucleosome footprints that are ready for library construction in less than 1 hour.

## Discussion

In this work, we have optimized and validated a rapid protocol for mapping nucleosome positions in eukaryotic genomes. While our protocol is simple and robust across organisms, there are caveats that should be noted prior to implementing the defined protocol. First, MNase activity can vary across lots and vendors, so it is critical to calibrate the MNase concentration to give the desired extent of digestion. We have successfully optimized MNase concentration from multiple vendors and strongly recommend initial enzyme titrations. Second, within an organism, the number of cells used or amount of MNase added could vary depending on the growth conditions or media composition. It is therefore possible that specific conditions or mutant strains may require subtle changes to digestion amount, particularly if it is challenging to accurately quantify the initial number of cells. Finally, there are likely organisms that may require additional initial steps to help permeabilize cells, particularly if the cells possess Zymolyase-resistant cell walls. Previous work has demonstrated success using cryogrinding as the cell breaking step instead of Zymolyase (Gonzalez and Scazzocchio 1997; Givens *et al.* 2011). The presented protocol is a reliable starting point from which adjustments can rapidly and easily be made.

In conclusion, this rapid MNase protocol is capable of accurately mapping nucleosome positions in *S. cerevisiae*, *S. pombe* and *N. crassa*, providing identical nucleosome positions as previously-published protocols in only 3 hours. In addition to recapitulating nucleosome footprints from model species in significantly less time, the same protocol can be used to isolate nucleosome footprints rapidly from wild fungal species for which no previous MNase protocol has been established. Finally, our rapid MNase protocol can successfully recover library-ready nucleosomal DNA from human cells in ~2.5 hours if cells are first crosslinked or in ~60 minutes if cells are not crosslinked. The single, simple protocol that rapidly recovers nucleosome footprints across the multiple tested organisms provides a unified strategy to map nucleosome positions. Because all steps are carried out in a single 1.5 ml tube using small volumes of reagents, it is likely that the described rapid MNase protocol can be easily adapted for automation. Based on our success in multiple systems, we believe that this simple protocol can be a first approach for mapping nucleosomes in organisms with or without established protocols.

## Materials and Methods

### Growth Conditions

For rapid MNase, an overnight culture grown in YPD was diluted to OD_600_ = 0.3 in 25 ml fresh YPD and grown at 30°C with shaking to a final OD_600_ = 0.8 (roughly 3*10^8^ cells). For “standard” protocol (Rodriguez et al. 2014), an overnight culture was diluted to OD_600_ = 0.1 in 250 ml of culture and grown to OD_600_ = 0.4-0.6. The culture was crosslinked with 1% formaldehyde (final concentration), and incubated at 30°C with shaking for an additional 15 minutes. Crosslinking was quenched by the addition of glycine (final concentration 125mM), and cells were pelleted at 3000xg for 5 minutes at 4°C. The pellet was resuspended in 1 ml of water, transferred to a 1.5 ml microcentrifuge tube, and pelleted at maximum speed for 20 s. Supernatant was removed completely with a pipette, and the pellet was flash frozen in liquid nitrogen and stored at −80°C.

For quiescent yeast, a 25 ml culture was grown at 30°C with shaking for 3 days in YPD and quiescent cells were purified by Percoll gradient as previously described (Allen et al. 2006; McKnight et al. 2015). The dense fraction was collected, pelleted, and OD_600_ was determined after cells were resuspended in water. A total of 100 OD_600_ units of cells (approximately 3*10^8^ cells) was crosslinked with 1% formaldehyde (final concentration) for 15 minutes, quenched with 125mM glycine, pelleted and flash-frozen in a 1.5 ml tube.

For yeast patches on YPD agar, a ~1 cm x 3 cm patch (Figure 2B) was made from a glycerol stock on YPD agar with appropriate selection and grown at 30°C overnight. The next morning the patch was collected with an inoculation loop and resuspended in 1 ml YPD. The OD_600_ of a 1:20 dilution was measured and 30 OD_600_ units of cells (roughly 10^8^ cells) was pelleted, resuspended in 1 ml PBS and crosslinked with 1% formaldehyde for 15 minutes. Crosslinking was quenched with 125mM glycine, then cells were pelleted and used directly in the rapid MNase protocol.

For *S. pombe*, an overnight culture was grown in YES media and diluted to OD_600_ = 0.3 in 25 ml fresh YES media. Cells were grown at 30°C with shaking until a final OD_600_ of 0.8 was reached. Cells were crosslinked with 1% formaldehyde for 15 minutes, quenched with 125mM glycine, pelleted and flash-frozen or used directly in the rapid MNase protocol.

For *N. crassa*, a 50 ml culture in Vogel’s medium with 1.5% sucrose was inoculated with 2.5*10^7^ cells, harvested from freshly grown conidia, and grown at 32°C with shaking for ~3.5 hours, until at least 70% of conidia had germinated as determined microscopically. Cells were crosslinked by addition of fresh formaldehyde for a final concentration of 1%, then shaken at room temperature for 10 minutes. The crosslinking reaction was quenched by the addition of 6.8 ml of 1M Tris-HCl pH 8 to a final concentration of 125mM and shaken for 10 minutes at room temperature. Germinated conidia were then harvested by centrifugation. Supernatant was carefully removed with a pipette, leaving some media with the pellet. Conidia and remaining media were transferred a 1.5 ml microcentrifuge tube and pelleted for 3 minutes. The pellet was resuspended in 1 ml of 1M sorbitol, pelleted again at 8000 rpm for 3 minutes and the final pellet was resuspended in ~200 ul of 1M sorbitol and stored at −80°C.

Wild mushrooms were picked on the University of Oregon campus and processed immediately or harvested in a nearby location and transported on dry ice prior to processing. Between 25 and 100 mg of fruiting body was excised and crosslinked with 1% formaldehyde in 1 ml phosphate-buffered saline for 15 minutes. Crosslinking was quenched with 125mM glycine, lightly disrupted with a wooden applicator stick, and tissue was pelleted at full speed in a microcentrifuge for 1 minute. The pellet was used directly in the rapid MNase protocol. Optimized samples (Figure 4B) used 50 mg of mushroom tissue.

PLB-985 cells (Tucker et al. 1987) were cultured to a density between 2*10^5^ and 2*10^6^ cells per ml in complete media [RPMI 1640 with HEPES and GlutaMAX (Gibco #72400120), supplemented with 10% heat-inactivated Fetal Bovine Serum (FBS), 100 IU penicillin, 100 ug/ml streptomycin] at 37°C in a humidified atmosphere with 5% CO_2_. One million cells were pelleted, resuspended in 1 ml PBS, crosslinked with 1% formaldehyde for 15 minutes while rotating, quenched with 125mM glycine, pelleted, and used directly in the rapid MNase protocol. For samples that were not crosslinked, 1 million cells were washed once with PBS and pelleted cells were used directly in the rapid MNase protocol.

### Micrococcal Nuclease Digestion of Samples

For rapid MNase, cells (*S. cerevisiae* pellets from liquid culture, *N. crassa*, *S. pombe* and wild mushrooms) were treated with 2 mg of Zymolyase (100T, AMS Bio) in a 1 ml solution of spheroplast buffer (1M sorbitol, 50mM Tris pH 7.5, 10mM beta-Mercaptoethanol) for 15 minutes while rotating at room temperature. Quiescent yeast were treated with 10 mg Zymolyase (100T, AMS Bio) for 1 hour. *S. cerevisiae* patches were treated with 2 mg Zymolyase (100T, AMS bio) for 30 minutes. PLB-985 cells did not require Zymolyase treatment. Cells were pelleted at full speed and resuspended in 100 ul MNase digestion buffer (1M sorbitol, 50mM NaCl, 10mM Tris pH 7.5, 5mM MgCl_2_, 1mM CaCl_2_, 0.075% (v/v) IGEPAL (Sigma), 0.5mM spermidine and 1mM beta-Mercaptoethanol; solution is stored at −80°C before use). Exonuclease III (NEB) (30 units) and micrococcal nuclease (Worthington) (60 units for *S. cerevisiae* pellets from liquid cultures, *N. crassa* samples, *S. pombe* samples, wild mushrooms, PLB-985 cells; 30 units for *S. cerevisiae* patches; 4 units for quiescent yeast) were added to samples and rapidly mixed. Digestion proceeded for 10 minutes at 37°C (or for indicated times for quiescent cells) and was quenched with a final concentration of 5mM EGTA and 5mM EDTA (from 50mM/50mM stock). RNase A (50 mg) was added for 30 minutes (15 minutes for uncrosslinked PLB-985 cells) at 42°C. Proteinase K (200 mg) was added for 45 minutes at 65°C (15 minutes for uncrosslinked PLB-985 cells). DNA footprints were recovered by MinElute PCR purification kit (Qiagen) and eluted with 12 ul of 1x Cutsmart buffer (NEB). For wild mushroom samples, after addition of buffer PB (Qiagen), samples were centrifuged for 2 minutes at full speed and the supernatant was applied to the MinElute column. Samples were treated with 5 units of Quick CIP (NEB) for 10 minutes at 37°C. DNA was separated by 2% agarose gel electrophoresis and mononucleosome bands were gel-extracted (Qiagen MinElute) and converted to genomic libraries using the Ovation Ultralow Library System V2 (NuGEN) according to the manufacturer’s instructions. Alternatively, Quick CIP was heat-inactivated and DNA was directly converted to MNase-seq libraries using the Ovation Ultralow Library System or similar high-throughput DNA library protocol with similar results.

For the “standard” protocol (Rodriguez et al. 2014), cells were resuspended in 40 ml of spheroplast buffer (see above recipe) and treated with 5 mg Zymolyase (100T, AMS Bio) for 15 minutes at room temperature and pelleted by centrifugation for 20 minutes at 4000 x g, 4°C. Spheroplasts were resuspended in 2 ml MNase digestion buffer (see above recipe) and divided into three 600 ul aliquots. Samples were digested with 10, 20 or 40 units of MNase (Worthington) and 30 units of Exonuclease III (NEB) for 10 minutes. Digestion was quenched with 1% SDS and 5mM EDTA (from 5%/50mM stock) and treated with 200 mg Proteinase K overnight at 65°C. DNA was then purified by phenol/chloroform extraction and ethanol precipitation. Recovered DNA was resuspended in 60 ul 1x NEB Buffer 2 and treated with 10 mg RNase A for 1 hour at 37°C. A 5 ul sample was analyzed by electrophoresis on 2% agarose gel and appropriately-digested samples were first purified by PCR purification kit (Qiagen), eluted with 50 ul of 1x NEB Buffer 3 and treated with 10 units of CIP (NEB) for 1 hour at 37°C. Phosphatase-treated DNA was purified with a MinElute PCR purification kit and eluted in 12 ul buffer EB. The full sample was separated by electrophoresis on 2% agarose gel and the mononucleosome band was excised, gel extracted (Qiagen MinElute) and used for library creation using the Ovation Ultralow Library System V2 (NuGEN). Libraries were sequenced at the University of Oregon’s Genomics and Cell Characterization Core Facility on an Illumina NextSeq 500 on the 37 cycle, paired-end, High Output setting, yielding approximately 20 million reads per sample.

### Data Processing

MNase sequencing data were analyzed as described previously (McKnight and Tsukiyama 2015). Briefly, paired-end reads were aligned to the *S. cerevisiae* sacCer3 (Yates et al. 2019), *S. pombe* (Wood et al. 2002) or *N. crassa* (Galagan et al. 2003) reference genome with Bowtie 2 (Langmead and Salzberg 2012), and filtered computationally for unique fragments between 100 and 200 bp. Dyad positions were calculated as the midpoint of paired reads, then dyad coverage was normalized across the genome for an average read/bp of 1.0. Nucleosome alignment to the Ume6 binding site, WNGGCGGCWW, was performed by taking average dyad signal at each position relative to all intergenic instances of a motif center. Intergenic instances of the Ume6 motif were found using the *Saccharomyces* Genome Database Pattern Matching tool (https://www.yeastgenome.org/nph-patmatch). Transcription start sites were obtained from published data sets for *S. cerevisiae* (Nagalakshmi et al. 2008) and *S. pombe* (Thodberg et al. 2019). Previously-published *S. cerevisiae* data (SRX5086833, (Donovan et al. 2019); SRX1176421, (McKnight et al. 2016)), *S. pombe* data (SRX554384, (Steglich et al. 2015); SRX331943, (DeGennaro et al. 2013)) and *N. crassa* data (SRX1596291, (Seymour et al. 2016); SRX2822417, (Klocko et al. 2019)) were downloaded from the Sequence Read Archive (SRA) and analyzed using our computational pipeline to identify nucleosome dyad positions. Data were visualized using Integrated Genome Browser (Freese et al. 2016). Sequencing data from this work can be accessed at the GEO database under accession code GSE141676.

## Author Contributions

Conceptualization, J.G.C. and J.N.M.; Methodology, L.E.M., J.G.C., T.B.B., V.N.T., and J.N.M.; Investigation, L.E.M., T.B.B., O.G.B.B., K.N.O., V.N.T., E.T.W., G.L.W., J.N.M.; Resources, G.L.W., E.T.W., D.A.D., S.D.H., E.U.S.; Writing - Original Draft, L.E.M., J.N.M.; Writing - Reviewing and Editing, L.E.M., J.G.C., T.B.B., O.G.B.B., K.N.O., V.N.T. E.T.W., G.L.W., D.A.D., S.D.H., E.U.S., J.N.M.; Visualization, J.N.M.; Supervision, S.D.H., E.U.S., J.N.M.; Project Administration, L.E.M., J.N.M.; Funding Acquisition, J.N.M.

## Data Availability

The datasets generated during this study are available at GEO under accession code GSE141676.

## Acknowledgments

The authors thank the Genomics Core (GC3F) at the University of Oregon for high throughput sequencing services. The authors also thank Gabe Zentner (Indiana University), Christine Cucinotta (Fred Hutchinson Cancer Research Center) and Andrew Klocko (University of Colorado, Colorado Springs) for helpful discussions related to this work. The authors also acknowledge the Finkelstein Lab for the BioRxiv template. Mushroom identification was assisted by Andrew Kern (University of Oregon), Nicole Liachko (University of Washington), Richard Ebright (Rutgers University) and Anthoni Goodman (University of Alabama-Birmingham). This work was supported by a National Institutes of Health training grant T32 GM007759 (to O.G.B.B., K.N.O., G.L.W., and D.A.D.), by a National Institutes of Health training grant T32 GM007413 (to V.N.T.), by NIGMS GM093061 (to E.U.S.) and by NIGMS R01 GM129242 (to J.N.M.).

